# Distinct brain-wide neural dynamics predict social approach behavior

**DOI:** 10.1101/2025.07.09.663340

**Authors:** Imri Lifshitz, Netta Livneh, Maayan Moshkovitz, Abeer Karmi, Lilach Avitan

## Abstract

Social behavior is essential for animal survival and adaptation, requiring the integration of sensory cues to guide interactions with conspecifics. A key component of social behavior is approach, where animals actively move toward social partners to maintain group cohesion, establish affiliations, and coordinate actions. While a continuous stream of social information is encoded across sensory modalities, it remains unclear whether a distinct neural process underlies social approach. Here, we developed a novel assay in which a head-fixed, tail-free zebrafish interacts with a freely swimming conspecific, enabling precise quantification of social behavior alongside whole-brain functional imaging at cellular resolution. We demonstrate that zebrafish approach behavior is jointly shaped by spatial and temporal information from conspecifics rather than by these features acting independently. Social approach behavior is preceded by distinct brain-wide neural activity patterns emerging seconds before movement onset, characterized by increased activity in a small subset of forebrain neurons and decreased activity in midbrain and hindbrain neuronal populations. These activity patterns reliably predict upcoming approach movements from each of these regions separately. Moreover, the extent to which neural activity distinguishes approach from non-approach movements predicts individual differences in social behavior, directly linking neural dynamics to behavioral variability. Together, our findings reveal a neural mechanism underlying social approach behavior, highlighting how a distributed yet functionally coordinated network facilitates social interaction.

## Introduction

Group living is a widespread strategy among animals, rooted in a fundamental social affiliative drive that motivates individuals to approach conspecifics ^1,2,3^. This affiliative behavior supports critical social interactions essential for survival, reproduction, and resilience to stress^4,5,6,7,8^. While progress has been made in identifying brain regions and sensory modalities mediating this behavior, key questions remain unanswered ^9,10,11^. Specifically, it is unclear how visual cues from conspecifics are processed to regulate social distance, how brain-wide neural dynamics mediate the approach toward conspecifics, and whether these dynamics differ from those underlying non-social behavior.

Several challenges hamper a comprehensive understanding of the neural processes underlying approach behavior toward a conspecific. First, the circuits driving this behavior are distributed across multiple brain regions, and capturing whole-brain neural activity with high spatiotemporal resolution remains technically unfeasible in mammals ^12^. Second, the interactive and dynamic nature of social behaviors makes it difficult to isolate the influence of the self and the other on individual behavioral and neural responses. To address these challenges, we used the zebrafish as a model system. Its vertebrate brain organization and genetic homology to mammals make it well-suited for uncovering conserved neural processes^13,14^. Additionally, its optical transparency enables whole-brain imaging at cellular resolution, allowing precise tracking of neural activity underlying social behavior. Zebrafish also exhibit visually driven social behaviors early in development, and their discrete movements allow natural segmentation of the behavior, facilitating the study of dynamic social interactions ^15,16,17,18,19^. Together, these advantages establish the zebrafish as a powerful system for dissecting the neural mechanisms underlying social approach behavior.

By two weeks of age, zebrafish maintain a short distance from conspecifics by actively approaching them ^16,17,19^. This behavior is abolished in the dark ^19^, suggesting a strong reliance on vision, although mechanosensory ^20,21,22^ and olfactory ^23^ contributions have also been reported. The approach toward conspecifics is known to be influenced by various spatial factors, including the number of conspecifics ^16,24^, the visual load they impose on the retina ^19^, and their shape ^25^. However, temporal coordination also plays a critical role. Freely swimming zebrafish exhibit synchrony with movements of conspecifics ^15,18,22,26^, and attraction to their movement statistics, even in the absence of reciprocal interaction ^17,27,28^. This suggests that approach behavior is not solely dictated by spatial cues but also depends on the precise timing of movements. Yet, how these spatial and temporal features interact to drive approach behavior remains unclear.

Our understanding of the neural circuits underlying fish social behavior has primarily relied on two complementary approaches. One uses simplified, synthetic social cues combined with functional imaging to isolate how specific cue features are encoded in the brain ^25,28^. The other captures natural social behavior and relies on post hoc proxies of neural activity. Although approaching conspecifics is central to social interaction, the lack of experimental paradigms that capture social actions during whole-brain functional imaging has left the real-time neural dynamics of approach behavior largely unexplored. In addition, zebrafish, like other social species, exhibit individual variability in social behavior, with some individuals displaying strong affiliative tendencies while others engage minimally with conspecifics ^15,29^. Whether and how this variability is reflected in neural activity remains an open question.

Here, we combined whole-brain neural recording at single-cell resolution with a novel behavioral assay to identify neural signatures of social approach behavior. To preserve the natural complexity of social cues, we introduced a freely swimming conspecific to a head-fixed, tail-free focal fish. This novel assay not only reproduced spatial and temporal features previously observed in freely shoaling fish, but also uncovered fundamental new principles of the behavior and its underlying neural processes. Specifically, focal fish were more likely to elicit approach movements when temporally coupling their movements with those of conspecifics, highlighting the interplay between spatial positioning and temporal coordination. Distinct neural processes in the midbrain, hindbrain, and forebrain consistently preceded approach movements, enabling the decoding of upcoming movements toward a conspecific in each region separately. Additionally, the extent to which neural activity distinguished approach from non-approach movements in each region accounted for individual variability in social behavior. These findings provide a comprehensive understanding of the brain-wide neural mechanisms underlying social approach behavior.

## Results

### A novel experimental assay for simultaneous recording of social interaction and whole-brain neural activity

To overcome the challenge of recording neural activity from a fish that is engaged in social interaction with a real conspecific, we developed a novel social assay. In this assay, a 15-17 days post fertilization (dpf) head-fixed, tail-free focal fish was embedded in the center of an inner transparent plate, observing an age-matched, freely swimming conspecific placed in an outer plate (n=44 fish pairs). The physical separation between the two plates ensured that the responses of the focal fish to the conspecific were driven solely by visual information. The con-specific could swim freely within an arc spanning 230°of the visual field of the focal fish (extending 115°to each side). While the behavior of both fish was captured with a high-speed camera, we recorded brain-wide neural activity of the focal fish using volumetric two-photon microscopy (Figure 1A,B, Figure S1A, Supplementary Movies 1-4, see Methods). Each experiment lasted 30 minutes, beginning with 5 minutes of darkness, during which the focal fish could not see the conspecific.

**Figure 1.**
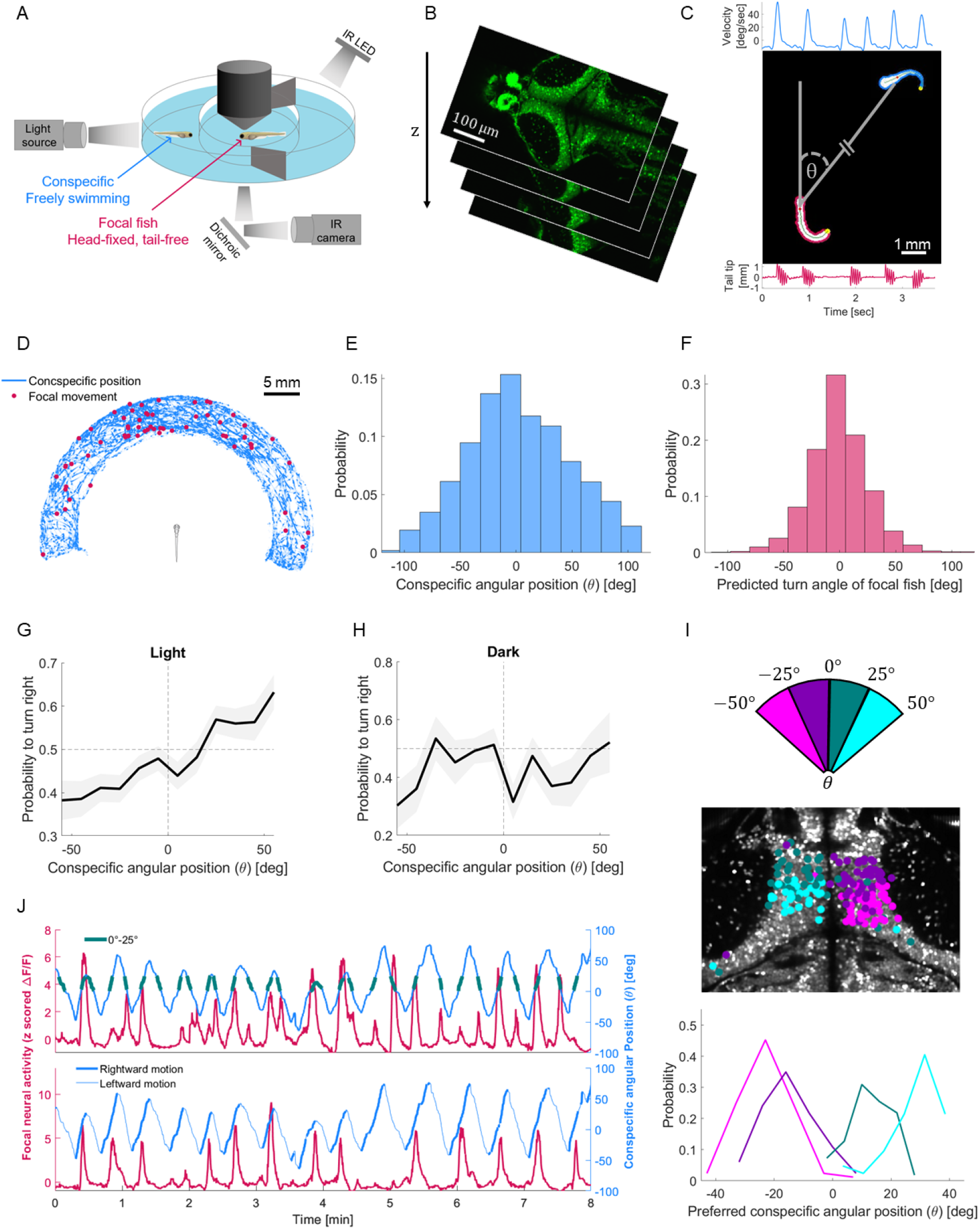
A novel social assay enables simultaneous recording of social behavior and whole-brain neural activity in cellular resolution. **A:** A schematic of the experimental social assay. A head-fixed and tail-free focal fish (red arrow) observes a freely swimming conspecific (blue arrow) while its whole brain neural activity is recorded using volumetric twophoton microscopy (8 planes, 10 µm apart, 3 volumes/sec). Behavior of both fish is captured at 166 Hz from below. **B:** Mean images of selected brain planes of the focal fish. **C:** Behavioral tracking of the focal fish (red contour) and conspecific (blue contour) enabled the segmentation of their movements, indicated by the angular velocity profile of the conspecific over three seconds (blue trace, top) and tail tip horizontal deflections of the focal fish over the same period (red trace, bottom). *θ* denotes the angular position of the conspecific relative to the heading direction of the focal fish. **D:** Swimming trajectory of a single conspecific during 30 minutes of experiment (blue). Red dots indicate the position of the conspecific at the time of focal fish movements, spanning the full range of angular positions. **E:** Distribution of the angular position (*θ*) of all conspecifics recorded, showing full-range coverage. Positive/negative angular positions correspond to the right/left sides of the focal fish, respectively. **F:** Distribution of the predicted turn angle of all focal fish movements, based on tracked tail dynamics (Figure S2). **G:** Probability of focal fish turning right as a function of the angular position of conspecifics in the light, calculated on all fish in 10°bins. Fish were more likely to turn right when conspecifics were on their right side and less likely to turn right when conspecifics were on their left. The gray shaded area indicates SEM across movements in each bin. **H:** Probability of focal fish turning right in darkness, when conspecifics were invisible, showing no tendency to approach conspecifics. **I:** Top: The frontal 100°visual field of the focal fish was divided into four 25°sections. Middle: Neurons responsive to the angular position of the conspecific in each section of the visual field were topographically organized. Bottom: Distributions of the preferred conspecific position of neurons corresponding to each section. **J:** Top: The conspecific angular position over eight minutes (blue trace), with green marks indicating times when the conspecific was within a specific visual section (0°-25°, corresponding to the green section in panel I). An example tectal neuron (red trace) selectively responded to this specific range of conspecific angular positions (0°-25°). Bottom: The same conspecific angular position over eight minutes as shown above (blue trace), with rightward motion indicated in dark blue. An example tectal neuron (red trace) selectively responded to rightward motion of the conspecific.

To rigorously quantify the behavior of both fish, we preprocessed the raw camera frames and tracked each fish separately using a custom-developed tracking system. For each fish, we extracted the contour and the tail midline from the swim bladder to the tail tip. These extracted features enabled tracking of the angular position and direction of motion of the conspecific relative to the focal fish, as well as its segments of movement throughout the experiment (Figure 1C, see Methods). Using the tail dynamics of the focal fish, we identified segments of movement and inferred the intended turn angle for each movement. This inference mechanism was trained and tested on recordings of freely swimming fish where tail dynamics and turn angles were measured and our prediction could be validated (Figure S2, see Methods). During the experiment, conspecifics spanned the entire arc area of the outer plate, and movements of the focal fish were elicited when the conspecific was at varying angular positions (Figure 1D). Across all fish, conspecifics spent 75% of their time between -50° to 50° relative to the focal fish (Figure 1E), where the predicted turn angle of 95% of focal fish movements fell within this range (Figure 1F). The movement rates of both the focal fish and conspecifics remained stable throughout the 25 minutes of recording in the light (Figure S1B). Together, this assay allowed accurate tracking of the fine details of two behaving fish where the focal fish observes a natural social cue.

Freely swimming 14 dpf larval zebrafish approach conspecifics by turning toward them, with a higher likelihood of turning right when conspecifics are in their right visual field and a lower likelihood when they are in their left ^16,19^. We examined whether this tendency is conserved in our assay by calculating the probability of the focal fish turning right as a function of the angular position of the conspecific. Consistent with previous results, fish turned toward their respective conspecifics, with approach probability increasing at larger angular positions (Figure 1G). The strength of this effect was comparable to that observed in freely swimming fish ^19^. Moreover, in line with previous findings, the tendency to approach a conspecific in our assay was abolished in the dark (Figure 1H) ^19^, and was independent of the direction of motion of conspecifics (Figure S1C)^30,31^. Hence, the behavior in our assay reproduced spatial findings previously observed in freely shoaling fish.

We next sought to verify that spatial information of the conspecific was encoded in tectal neural activity of the focal fish. To do so, we divided the visual field of the focal fish into four sections ranging from -50°to 50°, each covering 25° (Figure 1I, top). Using four regressors indicating the onset of conspecific movements within each section, we identified neurons whose activity correlated with the angular position of the conspecific (Figure 1J, top, see Methods). These neurons were topographically organized (Figure 1I, middle and bottom). In addition to the position of the conspecific, tectal neurons were correlated with its direction of motion (Figure 1J, bottom, Figure S1D). This confirms that the focal fish observed the natural social cue and encoded its physical properties, such as position and direction of motion, in tectal neural activity. Together, these findings support the use of our assay as a platform for studying the neural basis of social approach behavior.

### Approach behavior is jointly shaped by spatial and temporal information from conspecifics

By two weeks of age, freely shoaling larval zebrafish synchronize their movements by maintaining a short time lag relative to movements of conspecifics ^15,18,22^. We examined whether focal fish in our assay similarly relied on temporal information provided by their conspecifics to elicit movements. Focal fish did not elicit movements at random timings but tended to move shortly after conspecific movement onset, mostly during the glide phase, when angular velocity decreases (Figure 2A). To assess this relationship, we aligned the onset of each focal fish movement to time zero and examined conspecific behavior within a 1.2-second window before and after movement onset. Focal fish were most likely to move approximately 150 milliseconds after conspecific movement onset (Figure 2B, Figure S3A). This time lag between fish movements was significantly shorter than expected by chance (Figure 2C, Figure S3B). The empirical distribution of time lags between fish movements was further captured by a simple model of two Poisson processes when inducing a fraction of coupled movements, but not when the processes were fully independent, confirming that focal fish tended to synchronize their movements with those of the conspecifics (Figure S3C,D, see Methods). Together, fish in our assay replicated the synchrony observed in freely shoaling fish by temporally coupling their movements on a timescale of hundreds of milliseconds.

**Figure 2.**
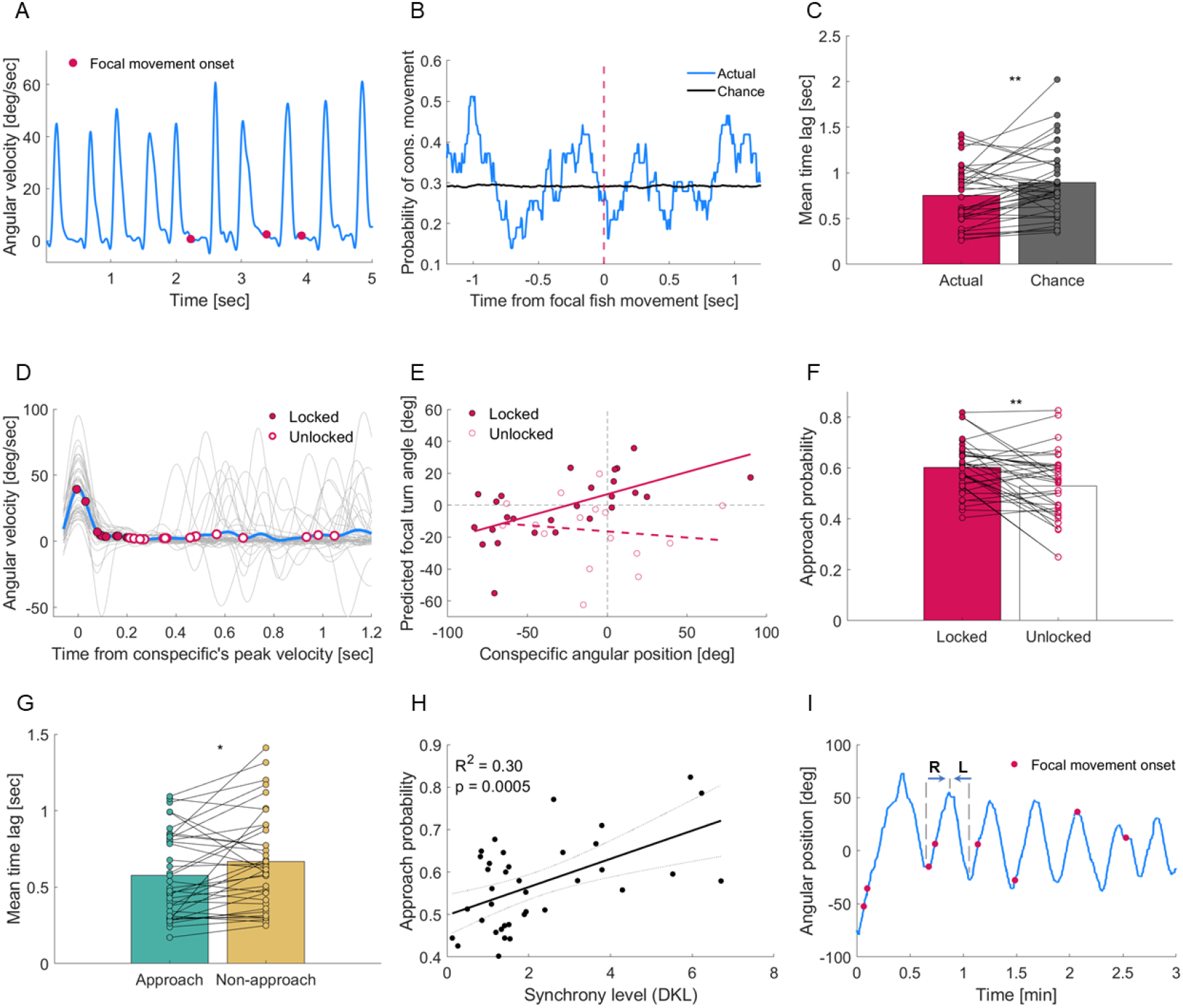
Spatial and temporal information from conspecifics interact to guide approach behavior. **A:** Angular velocity profile of an example conspecific (blue) over 5 seconds, with movement onsets of the focal fish overlaid (red dots), showing that focal fish movements typically occurred during the glide phase of conspecific movements. **B:** Probability of conspecific movements aligned to the onset of focal fish movement (red dashed line), showing that the focal fish maintained a short time lag relative to conspecific movements. Chance level probability (black line) was calculated based on the same number of focal fish movements elicited at random timings over 1000 iterations. **C:** The mean time lag between conspecific and focal fish movements was shorter in the actual data compared to the chance-level mean time lag, obtained by pairing each focal fish with a conspecific from a different experiment (paired t-test, *p* = 0.008). **D:** Angular velocity profiles of single conspecific movements (gray) that elicited a movement of the focal fish (red circles) and the mean conspecific velocity profile across all movements (blue). Filled and open circles indicate locked and unlocked movements, respectively, showing a concentration of locked movements at the glide phase of conspecific movement. **E:** The relationship between conspecific angular position and the predicted turn angle of the focal fish in an example pair, showing that locked movements were directed toward the conspecific, whereas unlocked movements were not (locked: *R*^2^ = 0.36, *p* = 0.001; unlocked: *R*^2^ = 0.02, *p* = 0.56; ANCOVA interaction effect: *p* = 0.027). **F:** Across fish, locked movements were more likely to be directed toward the conspecific than unlocked movements (paired t-test, *p* = 0.002). **G:** Approach movements exhibited shorter time lags compared to non-approach movements (paired t-test, *p* = 0.02). **H:** The level of synchrony between focal fish and conspecific movements, quantified as the distance between empirical and chance-level time lag distributions, was significantly correlated with approach probability. The chance-level distribution was obtained by using the same focal fish with a conspecific from a different experiment. The solid black line indicates the regression fit, with black dashed lines marking the 95% confidence bounds. **I:** The angular position of the conspecific over three minutes alternated between rightward and leftward movement sequences (blue). Example rightward and leftward sequences are indicated by blue arrows, with their duration bounded by gray dashed lines. Movement onsets of the focal fish are overlaid as red dots. Throughout the figure, ^*∗*^ and ^*∗∗*^ indicate *p <* 0.05 and *p <* 0.01, respectively, where specified.

Next, we asked whether spatial and temporal factors interact to modulate fish approach behavior. Specifically, we investigated whether focal fish movements occurring within the characteristic time lag following conspecific movements (locked movements) differed in their spatial relation to the conspecific compared to movements occurring outside this time lag (unlocked movements) (Figure 2D, see Methods). For each movement group, we examined the relationship between the angular position of the conspecific at movement onset and the predicted turn angle of the focal fish. We found a significant relationship for locked movements, but not for unlocked movements, with a significant interaction effect, suggesting that these relationships differ between movement types (Figure 2E). This indicates that movement synchrony is associated with an increased likelihood of turning toward a conspecific.

To further assess the relationship between synchrony and approach behavior across fish, we calculated the fraction of approach movements (defined as movements directed toward the conspecific, see Methods) for both locked and unlocked movements of each focal fish. Locked movements were significantly more likely to be approach movements than unlocked movements (Figure 2F). These locked movements were associated with higher conspecific movement frequencies, closer to the spontaneous frequency of 1.5 Hz at this age^17^ (Figure S3E). Furthermore, the mean time lag between focal fish and conspecific movements was shorter for approach movements than for non-approach movements (Figure 2G). The predicted turn angle of the focal fish did not differ between locked and unlocked movements or between approach and non-approach movements (Figure S3F), suggesting that different turn angles do not account for this spatio-temporal relationship. These results highlight that spatial and temporal information from conspecifics do not act independently but interact to guide approach behavior.

Freely swimming larval zebrafish exhibit considerable variability in their social behavior, with approximately 30% of 2-3-week-old larvae showing little to no social interaction ^15,29^. Consistent with these reports, 35% of fish in our dataset displayed low approach probabilities (Figure S3G). We asked whether this variability in fish approach behavior was related to their tendency to synchronize movements. To quantify synchrony level of each focal fish with its conspecific, we measured the distance between the empirical time lag distribution and a chance level time lag distribution (using Kullback–Leibler divergence, *D*_KL_, see Methods), with a larger distance indicating a stronger tendency to synchronize movements. We found that synchrony level varied across individuals and was correlated with approach probability (Figure 2H), suggesting that individual differences in movement synchrony contribute to variability in social approach behavior.

In addition to short-timescale synchrony, we examined whether focal fish movements also relate to longer timescale structures in conspecific behavior. Larval zebrafish have been shown to swim in alternating sequences of rightward and leftward turns, lasting up to 20 seconds^32^. Similarly, conspecifics in our assay exhibited rightward and leftward movement sequences relative to the focal fish (median duration: 9.5 seconds), with each sequence beginning when the conspecific changed its direction of motion (Figure 2I, Figure S3H). Focal fish tended to move early in these sequences (Figure 2I, Figure S3I), with approximately 60% of their movements occurring within the first half of a conspecific sequence. This suggests an additional temporal relationship, operating on a longer timescale, that links focal fish movements to changes in the motion direction of conspecifics.

### Distinct neural processes predict upcoming approach movements on a single-trial level

The established social assay enabled us to record activity from over 10,000 neurons across the brain of the focal fish during social interaction (Figure 3A, see Methods). While neural encoding of various sensory features of conspecifics, such as shape ^25^ and swim frequency ^28^ was studied, we examined whether distinct neural processes are associated with approach behavior toward conspecifics. We first characterized the neural activity preceding and following focal fish movements, using two regressors representing activity before and after each movement onset (Figure S4A, see Methods). Thousands of neurons across the brain showed increased activity following focal fish movements, whereas an order of magnitude fewer neurons displayed ramping activity prior to the movement which decreased afterward (Figure 3B). These two activity profiles were also captured by the first two principal components of the data (Figure S4B), highlighting the prominence of neural processes occurring before and after focal fish movements.

**Figure 3.**
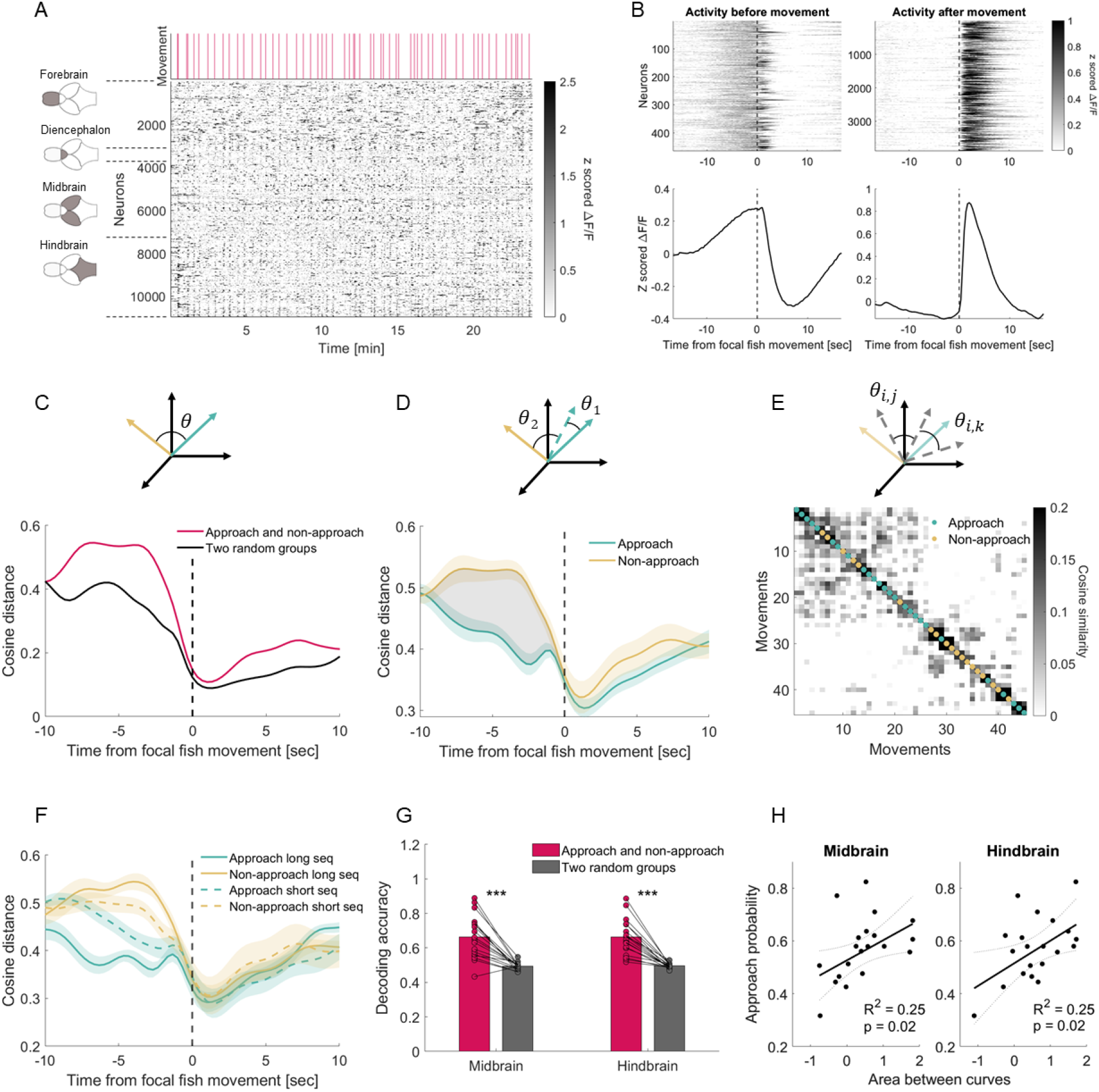
Distinct neural signatures predict upcoming approach movements and explain behavioral differences. **A:** A raster plot of 10,978 simultaneously recorded neurons from an example focal fish with the timing of its movements (red lines). Neurons are grouped according to their anatomical division into four major brain regions: forebrain, diencephalon, midbrain, and hindbrain (see Methods). **B:** Mean activity across movements of the focal fish for neurons that were active before (left) or after (right) the movements (top), with mean activity profile across these neurons (bottom). The dashed black lines at time zero indicate movement onset. **C:** Top: An illustration showing midbrain mean population vectors across approach (green) and non-approach (yellow) movements at a given time point. The angle between the vectors (*θ*) was used to calculate the cosine distance between them. Bottom: Cosine distance dynamics between mean population vectors of approach and non-approach movements over a 10-second window before and after movement onset (red curve), compared to the distance between the mean vectors of two randomly divided movement groups (black curve). Higher distances between approach and non-approach suggest distinct neural activity before these two movement types. The dashed black line at time zero represents movement onset. **D:** Top: An illustration of the angle between a population vector of a single approach movement (dashed green arrow) and the mean vectors across all approach (solid green) and non-approach (solid yellow) movements (excluding the tested single movement). Bottom: Cosine distance dynamics between any given population vector of a single approach movement and the mean vectors of approach (green) and non-approach (yellow) movements. Neural activity before approach movements showed greater similarity to the mean approach vector than to the mean non-approach vector. Green and yellow shaded areas indicate SEM across all tested movements in this example fish. Gray shaded area indicates the neural distinction between approach and non-approach movements before movement onset. Black dashed line at time zero represents movement onset. **E:** Top: An illustration of the angles between all pairs of population vectors (dashed gray arrows represent single movements). Bottom: Cosine similarity matrix between all pairs of vectors, where each vector represents the average neural activity over five seconds preceding a movement. The similarity matrix was hierarchically clustered to maximize the similarity between adjacent vectors, revealing two distinct clusters corresponding to approach and non-approach movements (green and yellow dots, respectively). **F:** The distinction in neural activity before approach and non-approach movements was associated with the time between the onset of conspecific sequence to the onset of focal fish movement. For long sequence durations (top 50%, solid lines), the distinction emerged earlier, while for short sequence durations (bottom 50%, dashed lines), it appeared closer to movement onset. **G:** Decoding accuracy of movement identity from preceding midbrain and hindbrain neural activity exceeded chance level (Bonferroni-corrected paired t-test, midbrain: *p* = 2 × 10^*−*5^, hindbrain: *p* = 4× 10^*−*6^). **H:** The area between the green and yellow curves shown in panel D for midbrain and hindbrain neural activity explained inter-individual variability in approach probability. Larger areas (indicating greater neural distinction) correlated with higher rates of approach movements. The solid black line indicates the robust regression fit, with black dashed lines marking the 95% confidence bounds. Throughout the figure, ^*∗∗∗*^ indicates *p <* 0.001, where specified.

We next examined whether neural activity linked to approach movements of the focal fish differs from that associated with non-approach movements, and whether such differences, if present, emerge before or after the movement. To address these questions, we first focused on the midbrain, a region known to be involved in fish social behavior ^13,25^. We calculated the mean population neural activity for approach and non-approach movements at each time point within a 10-second window before and after the movement. We then computed the cosine distance between the corresponding activity patterns (Figure 3C, top, see Methods). To control for baseline differences between arbitrary divisions into two movement groups, we repeated this analysis for two randomly selected groups. Distance values were higher before movement onset than after, both for approach vs. non-approach movements and in the control comparison (Figure 3C, bottom). This suggests a general phenomenon where neural activity diverges before fish movements and becomes more similar after movements. Interestingly, before movement onset, the distance between approach and non-approach movements exceeded that of the control groups, with this neural distinction emerging several seconds before the movement. Across fish, this distinction was evident in the midbrain and hindbrain but not in the forebrain and diencephalon (Figure S4C), suggesting that distinct neural processes in these regions precede approach movements toward a conspecific.

We then asked whether distinct neural processes were evident at the level of single movements. To test this, we first measured the distance between the population activity of each single movement of the focal fish and the mean population activity of both approach and non-approach movements (calculated on a leave-one-out basis, Figure 3D, top). Population neural activity of single approach movements more closely resembled the mean activity of approach movements than that of non-approach movements, with this distinction evident only before the movement (Figure 3D, bottom, Figure S4D). To further examine whether single approach movements are associated with similar neural patterns, we averaged the population neural activity over five seconds preceding each movement and computed pairwise similarity between all pairs of population vectors (Figure 3E, top), generating a similarity matrix. Hierarchical clustering of this matrix revealed two distinct clusters that largely corresponded to approach and non-approach movements (Figure 3E, bottom). Notably, these clusters did not correspond to right and left turns (Figure S4E), indicating that they reflect approach behavior rather than turning direction. Together, these results suggest that individual approach movements share a common neural signature emerging a few seconds before movement onset.

To examine whether the prolonged neural distinction emerging seconds before approach and non-approach movements reflects the long-timescale structure of conspecific behavior, we asked whether the emergence of this distinction depends on the duration of conspecific movement sequences. Conspecifics often moved in sequences of rightward and leftward movements (Figure 2I), with substantial variability in the time between the onset of such sequences and a focal fish movement (Figure S4F). We divided both approach and non-approach movements into two groups based on whether they followed a long or short sequence of conspecific movements (median threshold of 7.6 seconds between short and long sequences across fish, see Methods). In long sequences, distinct neural processes for approach and non-approach movements emerged approximately 10 seconds before movement onset, whereas in short sequences, this divergence emerged closer to movement onset (Figure 3F, Figure S4G). This suggests that the observed distinction in neural activity preceding approach and non-approach movements is related to the time elapsed following a change in the direction of motion of conspecifics.

Given the neural distinction before approach and non-approach movements, we asked whether the upcoming movement identity (approach or non-approach) could be decoded from the preceding population activity. We identified the neurons that contributed most to the distance between population responses preceding approach and non-approach movements (see Methods), focusing on the midbrain and hindbrain, regions where this distance was significantly evident across fish (Figure S4C). These neurons were spatially distributed throughout the midbrain and hindbrain (Figure S4H). Based on the mean activity of these neurons over the five seconds before movement onset, we decoded movement identity with accuracy significantly above chance in both regions (Figure 3G, see Methods). This confirms that fish approach behavior is represented by distinct neural activity in the midbrain and hindbrain and can be accurately predicted several seconds before movement onset.

A notable inter-individual variability in decoding performance was observed (Figure 3G), and while some fish showed higher performance than chance, others did not, presumably because the behavior was not fully established in all fish at this age (35% non-social fish). Similarly, the neural distinction preceding approach and non-approach movements (represented by the area between the curves, see Figure 3D) was larger than chance and showed considerable inter-individual variability (Figure S4I). Therefore, we asked whether these neural differences between individuals can explain the observed variability in approach behavior. Indeed, we identified a correlation between the area between the curves and the approach probability across fish (Figure 3H). This effect was not related to a bias in the fraction of right vs. left turns between approach and non-approach movements (Figure S4J). Taken together, these findings indicate that distinct neural dynamics in the midbrain and hindbrain predict approach behavior and account for its inter-individual variability.

### Opposing forebrain and midbrain-hindrain neural dynamics underlie socially specific approach behavior

Our results demonstrate that upcoming approach and non-approach movements can be predicted from midbrain and hindbrain neural activity preceding the movement based on the geometric distance between the respective population responses. We asked what pattern of neural activity determines this geometric distance and drives the distinction between these responses. To address this question, we examined the neurons that contributed most to this distance, and found that they exhibited reduced activity before approach movements (Figure 4A, top) and increased activity before non-approach movements (Figure 4A, bottom). Across fish, this distinction was evident before movement onset and gradually diminished after the movement (Figure 4B). To confirm that this neural pattern was related to the visual cue, we repeated the analysis using the same identified neurons for approach and non-approach movements elicited in the dark. Without visual access to conspecifics, the neural differences between approach and non-approach movements disappeared (Figure S5A), suggesting that the distinct neural activity preceding approach behavior in the midbrain and hindbrain arises specifically in response to the visual cue.

**Figure 4.**
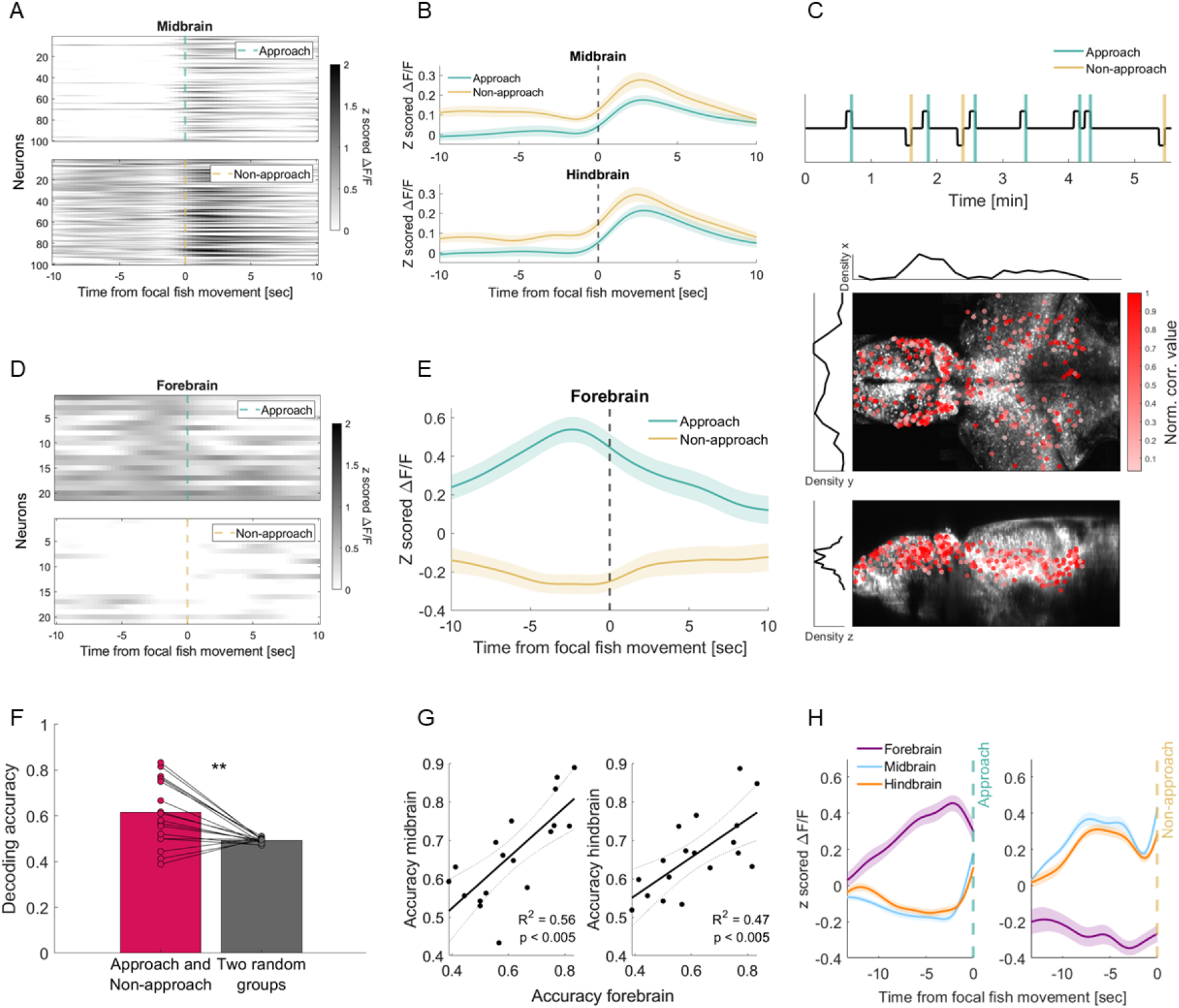
Concerted neural dynamics across the forebrain, midbrain, and hindbrain predict social approach behavior. **A:** Activity profiles of 100 midbrain neurons that most contributed to the distance between population responses preceding approach and non-approach movements in an example fish. These neurons showed reduced activity before approach movements (top) and increased activity before non-approach movements (bottom). Dashed green and yellow lines indicate the onset of approach and non-approach movements, respectively. **B:** Across fish, midbrain and hindbrain activity of the most contributing neurons was lower before approach movements than before non-approach movements. **C:** Top: “Approach regressor”, representing increased activity before approach movements and decreased activity before non-approach movements. Middle and Bottom: Neurons highly correlated with this regressor (399 neurons across 6 registered fish) were projected onto a reference brain, showing a dense concentration in the forebrain (shades of red indicate correlation strength). Top and side views are shown at the middle and bottom, respectively. **D:** Activity profiles of forebrain neurons highly correlated with the approach regressor in an example fish, showing higher activity before approach movements (top) and lower activity before non-approach movements (bottom). Dashed green and yellow lines indicate the onset of approach and non-approach movements, respectively. **E:** Across fish, forebrain neurons highly correlated with the approach regressor exhibited elevated activity before approach movements and reduced activity before non-approach movements. **F:** Decoding accuracy of movement identity from forebrain neural activity across the five seconds preceding movement was significantly above chance (paired t-test, *p* = 0.002). **G:** Decoding accuracy in the forebrain was correlated with accuracy in the midbrain and hindbrain. The solid black line indicates the regression fit, with black dashed lines marking the 95% confidence bounds. **H:** Coordinated brain-wide neural activity patterns in an example fish, showing elevated forebrain activity and reduced midbrain and hindbrain activity more than 10 seconds before approach movements, with the opposite pattern before non-approach movements. Throughout the figure, ^*∗∗*^ indicates *p <* 0.01, where specified. Shaded areas in Figures B, and E indicate SEM across fish, whereas in Figure H, they indicate SEM across neurons.

Since the neurons distinguishing approach from non-approach movements were suppressed before approach movements, we asked whether another set of neurons might be active, potentially responsible for their suppression. Therefore, we searched for neurons with elevated activity before approach movements and reduced activity before non-approach movements using an “approach regressor” obeying this pattern (Figure 4C, top, see Methods). We identified neurons across the entire brain that correlated with this activity pattern. Projecting the highly correlated neurons from different fish onto a reference brain (see Methods), we found a dense concentration of neurons in the forebrain, with fewer neurons sparsely distributed across other brain regions (Figure 4C, middle-bottom). These forebrain neurons demonstrated the expected pattern of elevated activity before approach movements and reduced activity before non-approach movements (Figure 4D,E). Similar to the midbrain and hindbrain, this distinction in forebrain neural activity was eliminated in the dark (Figure S5B), and was evident at the level of single movements (Figure S5C). These results suggest that approach movements are distinguished from non-approach movements also by the activity of a specific subset of forebrain neurons, with a pattern opposite to that observed in the midbrain and hindbrain.

The observed distinction in forebrain neural activity before approach and non-approach movements suggested that the identity of the upcoming movement could also be decoded from forebrain activity. To test this, we applied a logistic regression decoder, as previously used in the midbrain and hindbrain. For this analysis, we used the highly correlated neurons with the approach regressor (Figure 4C, see Methods). The decoder predicted the upcoming movement identity based on forebrain activity significantly better than chance (Figure 4F), with a higher effect size compared to other brain regions (Figure S5D). Interestingly, decoding performance in the forebrain correlated with the above-mentioned decoding in the midbrain and hindbrain (Figure 3G), with high decoding accuracy in the forebrain associated with high accuracy in the midbrain and hindbrain (Figure 4G), despite the use of distinct neuronal populations in each region. This highlights a coordinated brain-wide neural mechanism, with opposing activity patterns preceding approach behavior (Figure 4H). Moreover, similar to the midbrain and hindbrain, we found that greater neural distinction between approach and non-approach movements in the forebrain (measured by the area between cosine distance curves, Figure S5E) correlated with higher approach rates (Figure S5F). Together, these results show that distinct forebrain activity, which differentiates approach from non-approach movements, operates within a coordinated network that enables movement prediction and reflects individual differences in approach behavior.

Lastly, these findings raised the question of whether the observed neural distinction preceding approach behavior is specific to social cues or generalizes to any object the focal fish approaches. To address this question, we conducted an additional experiment with two counterbalanced sessions on the same focal fish (n=6). In one session, we recorded neural activity and behavior while the focal fish observed a real conspecific for 30 minutes. In another session of equal duration, the same focal fish observed a synthetic projected dot that followed the trajectory of a real conspecific but moved continuously (see Methods). On average across fish, approach probability did not differ between the two conditions (paired t-test, *p* =?0.09). However, distinct neural activity preceding approach vs. non-approach movements emerged only when focal fish observed a real conspecific (Figure S5G). Additionally, the opposing patterns - decreased midbrain and hindbrain activity alongside increased forebrain activity before approach movements - were observed only when fish approached a real conspecific, and were absent when they approached a moving dot (Figure S5H). These results confirm that the coordinated, brain-wide neural processes preceding approach behavior are not a general feature of approach movements but are specifically linked to social cues.

## Discussion

Social interactions are dynamic, complex, and high-dimensional, presenting major experimental and computational challenges for uncovering their behavioral principles and underlying neural mechanisms. We developed a novel assay for simultaneous recording of whole-brain neural activity and social interaction in zebrafish, revealing fundamental behavioral and neural principles of social approach behavior toward a conspecific. By preserving the full complexity of social cues, we found that approach movements are driven by the interplay of spatial and temporal features. These approach movements were preceded by distinct neural activity patterns in a small subset of forebrain neurons and neuronal populations in the midbrain and hindbrain, revealing a specific brain-wide signature emerging seconds before social approach. These distinct patterns enabled accurate decoding of upcoming social actions from each region separately, demonstrating that social information processing is distributed across circuits well before execution. Furthermore, the extent to which neural activity distinguished approach from non-approach movements predicted individual variability in social behavior, linking behavioral differences with the distinction of neural dynamics. By combining high-resolution whole-brain imaging, ethologically relevant social stimuli, and a novel assay to capture social actions, this work provides a detailed mechanistic and brain-wide account of the neural processes underlying social approach behavior at single-cell resolution.

Brain-wide, interconnected circuits regulating social behavior are an evolutionarily conserved feature across diverse taxa, from fish to birds and mammals ^13,14,33^. Consistent with this framework, our results reveal that social approach behavior is not mediated by a single brain region but instead is distributed across multiple areas. Our work extends this understanding by identifying distinct neural dynamics within specific forebrain, midbrain, and hindbrain populations, which exhibit opposing activity patterns before approach movements. Interestingly, forebrain activity was driven by a small subset of neurons (25 on average across all fish), compared to a few hundred neurons in the midbrain and hindbrain, which may explain why the overall forebrain population did not reveal the distinction between approach and non-approach movements (Figure S4C). These forebrain neurons were primarily located in the pallium (dorsal forebrain, Figure S5I), a region known to be involved in social behavior in fish ^34^. While the ventral forebrain has been implicated in various aspects of social behavior and its neural encoding ^23,35,36,37^, our findings support the role of the dorsal forebrain in mediating social engagement. Although the underlying connectivity of these identified pallial neurons is yet to be uncovered, forebrain neurons that synapse at the midbrain and hindbrain have already been reported ^38^. Further work is required to trace the connectivity of these neurons and uncover their neurotransmitter identity to account for these distinct and opposing neural patterns specific to approach behavior.

Our results uncovered prolonged neural dynamics that emerge several seconds before the onset of a social movement. This long-timescale activity aligns with previous reports of sustained neural signals preceding movement initiation, driven by global visual motion ^39,40^. These signals have been suggested to reflect evidence accumulation processes, and their temporal profile parallels primate models in which neural activity builds progressively until a behavioral threshold is reached ^41^. Consistent with these findings, our results indicate that social movements in zebrafish arise from long-timescale neural computations, linked to the time elapsed between a change in conspecific direction of motion and focal fish movement onset.

Our data reveal substantial inter-individual variability in approach behavior, consistent with previous reports ^15,29^. By analyzing this variability, we identified both behavioral and neural factors that may account for these differences. From a behavioral perspective, we found that focal fish with stronger temporal coupling between their movements and those of conspecifics were more likely to approach conspecifics (Figure 2H). This suggests that the ability to synchronize with social partners is a key component of social behavior. From a neural perspective, we observed that the more distinct the neural activity patterns preceding approach and non-approach movements, the higher the probability of approach behavior (Figure 3H, Figure S5F). This finding suggests that the formation of shoaling behavior may arise from a progressive refinement of these neural distinctions. Further research is required to uncover how this neural distinction between approach and non approach movements is shaped by development ^15,16,17,19,25^, prior social experience ^20,25,29,42^, neuropeptides ^21,23,27,36,37,43,44^, genetics ^45,46,47^, and internal states ^23,48^. Our work established a robust framework for dissecting the contribution of each factor at the functional neural level.

Social interactions range from simple, reflexive, and unidirectional behaviors to complex, reciprocal, and cognitively demanding exchanges. Despite this diversity, they share fundamental features: actions that are temporally coupled and spatially oriented with respect to a conspecific. These principles are evident across species, from collective odor responses in flies ^49^ to parental behavior in mice ^10^ and to gaze following in humans ^50^. In mammals, coordinated social actions rely on interactions between prefrontal circuits, subcortical reward systems, and motor areas^9^. Our findings indicate that zebrafish engage similar structures during social approach behavior despite their simpler neural architecture. Moreover, we identify a brain-wide signature preceding social approach movements, which is robust enough to predict whether an upcoming movement will be directed toward a social partner. Whether prefrontal activity modulates striatal and brainstem circuits using similar activity profiles remains an open question. Zebrafish offer a unique opportunity to study social behavior at the cellular level across the entire vertebrate brain, advancing our understanding of brain-wide neural dynamics mediating social behavior.

## Acknowledgements

We thank Ami Citri, Inbal Goshen, and Inbal Shainer for their comments on earlier versions of the article. In addition, We thank Raunak Basu, Shlomo Wagner, and the Avitan lab members for the inspiring and stimulating discussions. Finally, we would like to thank ELSC’s fabrication laboratory team, and in particular Itamar Frachtenberg, for invaluable assistance with building our setup.

## Methods

### Zebrafish maintenance

Nacre zebrafish (Danio rerio) embryos expressing elavl3:H2B-GCaMP6s^51^ were collected and raised according to established procedures ^52^ and raised at 27°C with a light and dark cycle of 14/10 hours. Fish were fed live rotifers (Brachionus plicatilis) daily from 5 dpf. Experiments took place when zerbrafish were 15-17 dpf. At this age, the sex of the fish cannot be determined. All procedures were performed with approval from The Hebrew University of Jerusalem Animal Ethics Committee. The Hebrew University is accredited by the Association for Assessment and Accreditation of Laboratory Animal Care International (AAALAC).

### A novel experimental social assay

A single focal fish was embedded head-fixed and tail-free in 2.5% low-melting agarose at the center of a 33 mm diameter plate. This plate was placed within a larger 58 mm plate, held in place by custom-made clips, with the space between the plates filled with fish media. A freely swimming conspecific was introduced into the outer plate, occupying 230°of the focal fish’s visual field at a distance range of 16.5-29 mm from the focal fish, aligning with the reported attraction zone ^17,20,28^. Behavioral recording of both fish was conducted from the bottom using a FLIR Grasshopper 3 camera at 166 Hz. Simultaneously, volumetric two-photon imaging (Nikon objective, CFI ×16, 0.8 NA) captured neural activity over a 705 × 352 × 70 µm area at 3 volumes/sec. A projector (Optoma ML750ST) illuminated the outer plate of the conspecific, and the plates were further illuminated using infrared LEDs (850 nm). The water temperature was set to 27°C, maintained with a thermoplate, and monitored using a thermocouple (Tokai Hit). The experiment began with a 5-minute dark phase, after which a physical shutter was removed to allow the illumination of the plates.

Focal fish that elicited movements almost exclusively to one side (absolute median predicted turn angle *>* 25°, n=3) were excluded from the dataset. Non-social fish, with approach probability below 0.4 (n=6/44), were excluded from the quantification of social behavior (Figures 1,2), without affecting the significance of any reported effect. Fish were excluded from the neural analysis if their recordings showed z-axis drift during functional imaging, exhibited low signal-to-noise ratio, or had synchronization issues between the microscope and the camera. This resulted in functional neuroimaging data from 21 fish. All details of the control experiment (Figure S6G,H) were identical, except that instead of interacting with a real conspecific, focal fish observed a synthetic moving dot occupying 4 degrees of the visual field. This dot followed the trajectory of an example real conspecific but moved continuously.

### z-Stack acquisition and image registration

Subsequent to the functional imaging, the entire brain structure (300-350 imaging planes, 1 µm apart) of several focal fish (n=6) was scanned for structure. While a brain atlas for 6 dpf zebrafish is available, no published atlas currently exists for 15–17 dpf fish. Therefore, these brains were registered to a reference brain (selected from the recorded brains for function and structure), aligning all neurons within the same coordinate system (Figure 4C, Figure S4H). The registration process was done using Advanced Normalization Tools (ANTs^53^). In the control experiment, the brain structure of the focal fish was scanned after the second session (conspecific/dot in a balanced order) and subsequently registered to the reference brain. In a separate process, we registered our 16 dpf reference brain onto the 6 dpf MapZebrain atlas ^38^ to identify the forebrain structure where the highly correlated neurons reside. Despite the difference in developmental stages, it allowed us to roughly locate the identified neurons in the dorsal part of the forebrain (Figure S5I).

### Preprocessing of neural data

The raw volumetric two-photon imaging data were processed using Suite2p ^54^, including motion correction and cell detection. The Δ*F/F* traces for each cell were then calculated, with the baseline set to the 8th percentile of a moving window. An additional denoising step excluded cells whose Δ*F/F* distribution was not different from a Gaussian, a characteristic of non-responsive noisy neurons. Similar to a previous study ^25^, the brain of each fish was manually divided into four major regions: forebrain, diencephalon, midbrain and hindbrain.

### Behavioral tracking and feature extraction

Fish behavior was tracked using a custom-made algorithm originally developed for tracking natural behavior in freely swimming fish ^55^. To remove stationary objects and maintain pixels that varied in their brightness, each raw frame was converted into a binary image by subtracting from each pixel its minimal intensity value recorded throughout the experiment. This step captured fish movements while eliminating stationary background objects. These binary images were then used as input for our algorithm, which tracked the behavior of each fish (focal and conspecific). This step resulted in the extraction of key features, including the fish contour and the tail midline, extending from the swim bladder to the tail tip. To detect the timing of movements for both fish, we examined the velocity of the points along the tail and set a threshold for mean velocity across points to identify the onset and offset of each movement.

### Predicting turn angle from tail dynamics

To develop a mechanism for predicting turn angles, we analyzed tail dynamics in freely swimming fish, where tail postures and the corresponding turn angles were accurately tracked and quantified. We used a natural behavior dataset in which individual fish (n=15) were recorded for 15 minutes in a plate containing paramecia, using high spatiotemporal resolution imaging (500 Hz, 0.02 mm/pixel, Figure S2A)^55^. During this period, fish engaged in both hunting behavior, detected by eye convergence, and exploratory behavior, characterized by diverged eyes. We tracked the tail midline from the swim bladder to the tail tip across all movements. Each tail posture was divided into 100 equidistant segments, and the angle of each segment relative to the horizon was computed (Figure S2B). A high-dimensional representation of movement could thus be defined by a series of segment angles throughout the movement (Figure S2C).

Previous studies have shown that tail postures are highly characteristic, with three principal components (PCs) explaining most of the variance in tail movements ^56,57^. We reproduced this result in our dataset, showing that in both freely swimming fish and head-fixed, tail-free fish, the first three PCs accounted for approximately 85% of the variance (Figure S2D). Importantly, the PCs extracted from head-fixed fish closely resembled those of freely swimming fish (Figure S2E), suggesting a strong similarity in tail postures between the two conditions. Each movement consisted of a sequence of tail postures, which could be represented as a time series of coefficients along the three PCs (Figure S2F). Since fish first turn and then move forward or backward ^55^, we focused on PC coefficients during the initial 30 ms of each movement, concatenating them into a single row vector. Tail dynamics were therefore represented in a matrix (**A**), where each row is a movement and each column is a frame. For freely swimming fish, we measured changes in turn angle, constructing the result vector 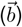. We then used a linear regression model to find a transformation vector 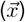 that would optimally convert PC coefficient vectors into the corresponding turn angles (Figure S2G).

Once the transformation vector was extracted, we evaluated its accuracy by comparing predicted and actual turn angles in freely swimming fish. We first trained the model on hunting movements of one fish and tested it on hunting movements of another fish. The model accurately predicted turn angles (Figure S2H), suggesting that it could be generalized across individuals. We then trained the model on hunting movements and tested it on exploratory movements from another fish, where it also accurately predicted turn angles (Figure S2I), indicating applicability across different behavioral contexts. More broadly, these findings indicate the existence of a robust and reproducible mechanism for interpreting tail dynamics and inferring turn angles. Since this transformation vector generalized across fish and behavioral contexts, and given that PCs from head-fixed fish closely resembled those from freely swimming fish, we applied this transformation vector to predict the turn angles of focal fish based on the PC coefficients of their movements.

### Assessing movement synchrony

The synchrony level of each focal fish was measured using the Kullback–Leibler divergence (*D*_KL_), between its empirical time lag distribution and a chance-level distribution, calculated using a conspecific from a different experiment. For each focal fish, the conspecific with the closest number of movements to the actual conspecific was selected. To account for the asymmetry of *D*_KL_, we averaged this measure over both directions of comparison. The observed synchrony in our dataset was also demonstrated using a Poisson process-based model with three parameters drawn from the empirical data: the movement rates of the focal fish and the conspecific, and the fraction of locked movements elicited by the focal fish (coupling fraction). The chance-level distribution of the model was generated using two independent processes with no synchrony.

### Classification of focal fish movements

The classification of locked and unlocked movements was based on the time lag distribution between the movements of each focal fish and those of its respective conspecific. A threshold was set for each fish based on the difference between the empirical time lag distribution and the chance-level distribution, allowing us to identify the value below which the empirical distribution exceeded the chance-level. Focal fish movements with time lags below or above this threshold were defined as locked or unlocked movements, respectively.

The classification of approach and non-approach movements was based on the predicted turn angle of the focal fish. Movements with a predicted angle greater than 5°to either side (right or left) were defined as approach movements if directed toward the half of the visual field where the conspecific was located (e.g., rightward movements when the conspecific was on the right). Movements directed to the opposite side of the conspecific were defined as non-approach. Analyses based on the probability or average of approach/locked movements included fish with more than eight movements of that type to allow robust statistics.

The division of approach and non-approach movements according to long and short conspecific sequences was based on the duration from the onset of the closest conspecific sequence to the movement onset of the focal fish. For each movement group (approach and non-approach), the median duration from conspecific sequence onset was used as a threshold: movements elicited after a shorter/longer duration than the median were classified as movements following a short/long sequence, respectively.

### Identifying neurons correlated with regressors

To identify tectal neurons correlated with the angular position of the conspecific, we divided the frontal 100°of the focal fish’s visual field into four sections. A separate regressor was generated for each section, with regressor activity levels set to one when the conspecific moved within the corresponding section of the visual field. Each regressor was convolved with the temporal response kernel of GCaMP6s. Neurons with correlation values exceeding two standard deviations above the mean of the distribution of all neurons were considered highly correlated. Next, for each neuron within these four populations, we identified peaks in activity and averaged the conspecific angular position across all peaks, allowing us to quantify the preferred angular position of each neuron (Figure 1I).

To identify neurons responsive to the direction of conspecific motion, we constructed two regressors (convolved with the GCaMP6s kernel) that were active at the onset of rightward or leftward conspecific sequences. Additionally, two regressors with the same number of sequences, but with randomly drawn timings, were constructed. Neurons were considered highly correlated if their correlation value with the actual regressor exceeded the 99th percentile of the corresponding random correlation value (Figure S1D). This cutoff was applied to the movements and approach regressors as detailed below.

To identify neurons active before or after focal fish movement, we constructed two box regressors with elevated activity either preceding or following each movement, both lasting 6.66 seconds (Figure S4A). Two additional control regressors were generated with the same number of movements, but with randomly drawn timings. The 99th percentile of the correlation values with the control regressors was set as a threshold.

Similarly, we constructed a box “approach regressor” (Figure 4C, top) with elevated activity before approach movements and reduced activity before non-approach movements. The corresponding control regressor exhibited elevated activity before a randomly selected group of focal fish movements and reduced activity before another group. The 99th percentile of the correlation values with the control regressor was set as a threshold.

### Cosine distance and decoding movement identity

The cosine similarity between two population vectors **v**_1_ and **v**_2_ was computed as cos 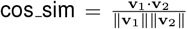, and the cosine distance was then calculated as 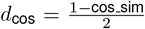. Dividing by 2 normalized this measure to a range of 0 to 1, where 0 indicates identical vectors and 1 indicates vectors pointing in opposite directions. To identify the neurons that contributed most to the distance between the mean population vectors of approach and non-approach movements, we selected the 100 neurons with the smallest average contribution to the dot product **v**_1_ · **v**_2_ in the 10 seconds preceding the movement. This analysis was done separately for each focal fish and brain region.

To prevent contamination from neural activity associated with preceding movements, all cosine distance analyses excluded movements separated by less than 10 seconds from the preceding movement.

To decode the upcoming movement identity (approach and non-approach) based on midbrain and hindbrain neural activity (Figure 3G), we used a logistic regression decoder with leave-one-out cross-validation. For each fish and brain region, the 300 most contributing neurons were selected in each iteration using a training set, and their mean activity over the five seconds before movement onset was used to predict the left-out movement. Using the same procedure, we decoded movement identity of the two control groups, repeating it 50 times with different random realizations. To decode movement identity based on forebrain neural activity (Figure 4F), we used the same logistic regression model, selecting the 20 neurons most highly correlated with the “approach regressor.” The number of movements in each group was balanced, and fish with more than 10 movements in each group were included.

### Statistical analysis

All statistical details are provided in the figures and their captions. All tests were two-tailed unless otherwise specified. For multiple comparisons, p-values were Bonferroni-corrected.

