## Supplementary info for "Distinct brain-wide neural dynamics predict social approach behavior"

### Supplementary information

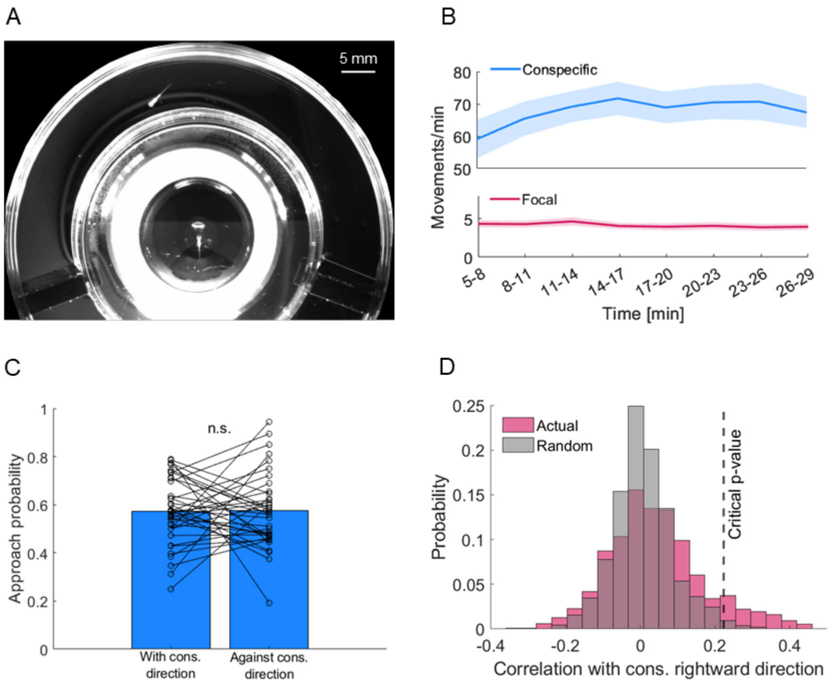


**Figure S1: Characteristics of the novel social assay. A:** A raw camera frame showing two behaving fish in the social assay. **B:** Movement rates of conspecifics (top, 68 movements/min on average) and focal fish (bottom, 4.1 movements/min on average) remained stable over 25 minutes in the light (Repeated measures ANOVA, focal fish: $p = 0.39$, conspecific: $p = 0.17$). Shaded areas indicate SEM across all fish. **C:** The fraction of approach movements directed with or against the direction of motion of conspecifics did not differ, suggesting no directional preference (paired t-test, $p = 0.94$). **D:** Distribution of correlation coefficients between neural activity of each tectal neuron and a regressor indicating conspecific rightward motion (red). The random correlation distribution was calculated using a regressor constructed from the same number of movements with randomly selected time points (gray). The critical p-value was the 99th percentile of the random distribution.

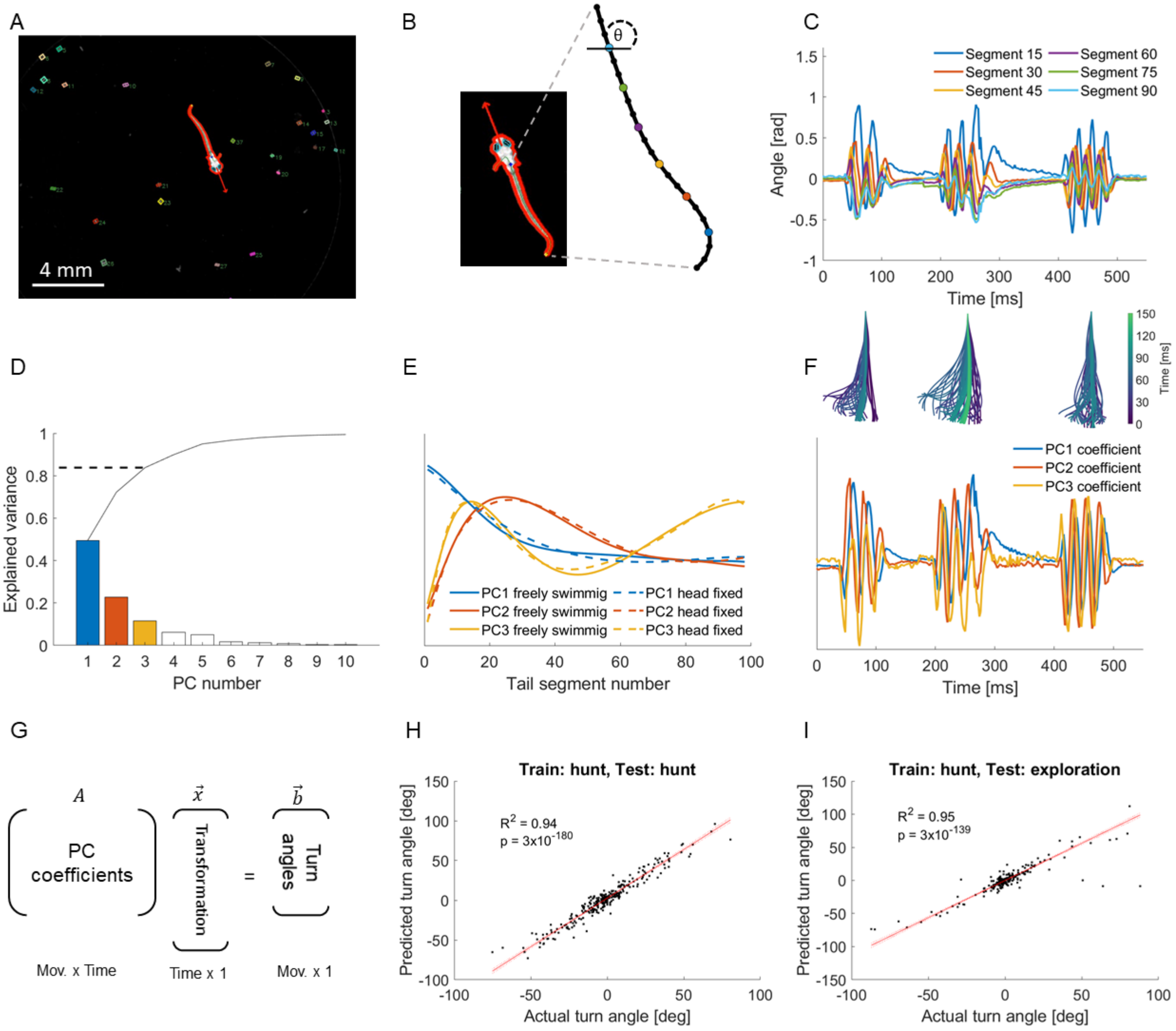


**Figure S2: Fish turn angle is accurately predicted from its tail dynamics.** **A**: An example frame of a freely swimming hunting zebrafish, tracked and annotated by our algorithm[1]. The algorithm identified the fish contour (red contour), heading direction (red arrow), swim bladder (blue point), tail midline (green line), tail tip (yellow point), and eyes (cyan). Paramecia (prey items) are marked by colored rectangles. This natural behavior dataset included 15 minutes of behavior in a plate containing paramecia, where hunting events were interleaved with segments of exploratory behavior[1]. **B:** Fish tail midline was divided into 100 equidistant segments, and the angle between each segment and the horizon ($\theta$) was extracted. **C:** The angle dynamics of selected tail segments across three movements showed an oscillatory pattern. Colors indicate different segments, corresponding to the colored points on the tail shown in panel B. **D:** Principal components (PCs) of tail postures were extracted using singular value decomposition (SVD). The first three PCs explained 87% of the variance of tail postures in freely swimming fish. **E:** The first three PCs of tail postures of an example freely swimming fish (19,952 postures, 570 movements) and an example head-fixed, tail free fish (1,166 postures, 31 movements, 84% explained variance by the first three PCs). These PCs were highly characteristic and similar in freely swimming fish and in head-fixed fish in our assay. **F:** Top: Tail postures over time for the same three movements as in panel C. The scale bar indicates time progression within each movement. Bottom: The same three movements are represented as the coefficients of the first three PCs over time. **G:** A transformation, mapping the first three PC coefficients of each movement to the actual turn angle, was obtained using a linear regression model. The transformation vector ($\vec{x}$) was then used to predict the intended turn angles of another fish ($\vec{b}$) based on the PC coefficients of its own tail movements ($\mathbf{A}$). **H:** Training the model on one hunting fish and testing on another hunting fish provided accurate predictions and indicated that this transformation can be generalized to other fish. Each dot represents a single movement. The solid red line indicates the robust regression fit, with red dashed lines marking the 95% confidence bounds. **I:** Training on one hunting fish and testing on another exploring fish yielded accurate predictions, indicating that the transformation generalizes across behavioral contexts.

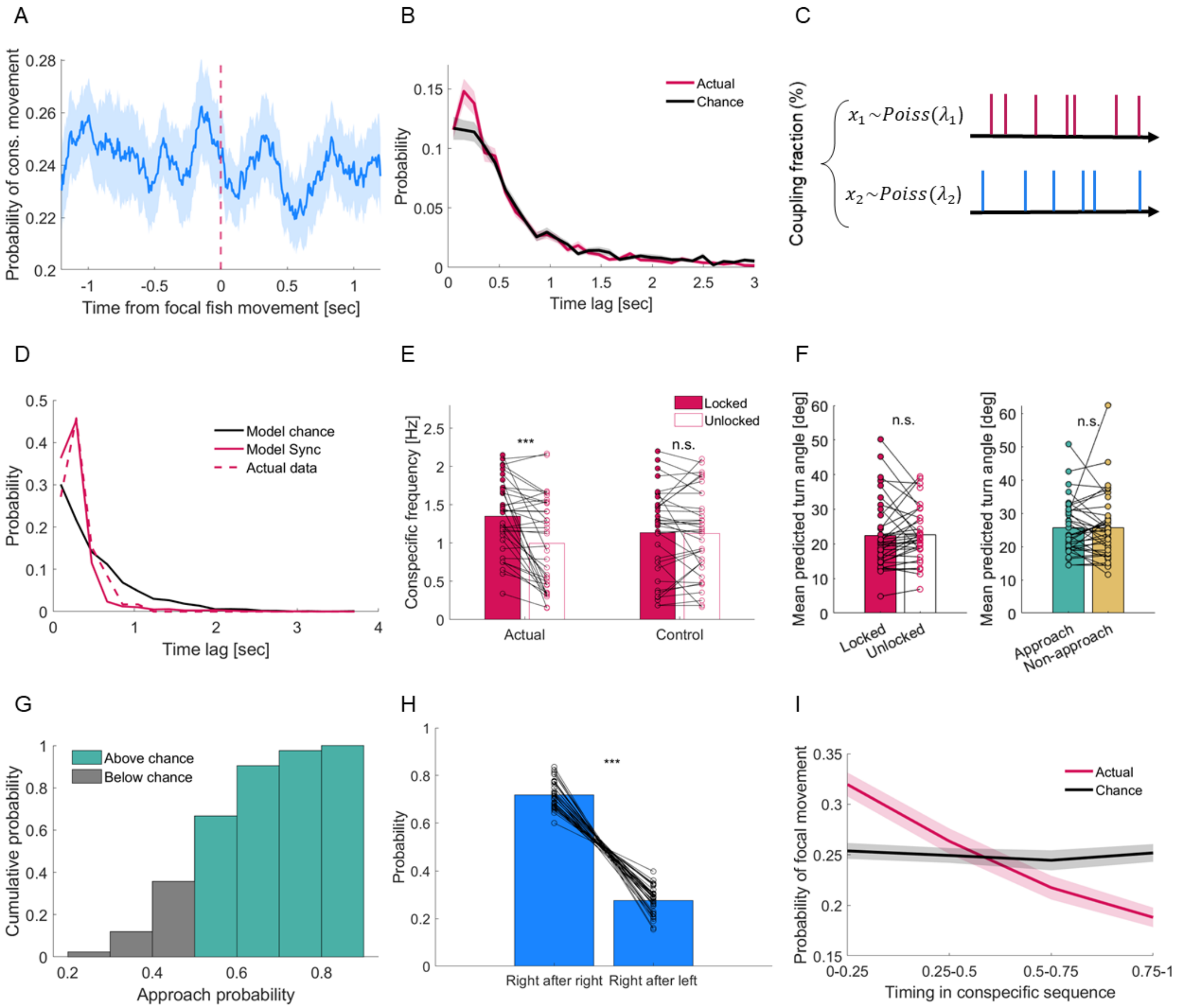
A
Probability of cons. movement
Time from focal fish movement [sec]
B
Probability
Time lag [sec]
Actual
Chance
C
Coupling fraction (%)
$x_1 \sim Poiss(\lambda_1)$
$x_2 \sim Poiss(\lambda_2)$
D
Probability
Time lag [sec]
Model chance
Model Sync
Actual data
E
Conspecific frequency [Hz]
Locked
Unlocked
***
n.s.
Actual
Control
F
Mean predicted turn angle [deg]
n.s.
Locked
Unlocked
Approach
Non-approach
G
Cumulative probability
Above chance
Below chance
Approach probability
H
Probability
***
Right after right
Right after left
I
Probability of focal movement
Actual
Chance
Timing in conspecific sequence
0-0.25
0.25-0.5
0.5-0.75
0.75-1

**Figure S3: Focal fish synchronize their movements with conspecifics on short and long timescales. A:** Probability of conspecific movements relative to focal fish movement onset, demonstrating that focal fish synchronized their movements with conspecifics within a short time lag. Shaded areas indicate SEM across all fish. **B:** Empirical distribution of time lags between conspecific movements and subsequent focal fish movements (red), overlaid on a chance-level distribution (black), calculated using a conspecific from a different experiment. The mean distributions differed significantly (KS test, $p = 0.01$). Shaded area indicate SEM across all fish. **C:** A Poisson process-based model of movement timing, using empirical movement rates for both focal fish and conspecifics, with a fraction of movements set to occur within a short time lag (sync). **D:** The sync model reproduced the time lag distribution of an example fish (with coupling fraction of 81%). The time lag distributions of the model and the data were not different (KS test, $p = 0.33$). The time lag distributions of the data and the sync model were both different from chance level time lag distribution (KS test, data vs. chance: $p = 9 \times 10^{-4}$, sync vs. chance: $p = 6 \times 10^{-174}$). p-values were Bonferroni-corrected. **E:** Locked movements were elicited when conspecifics moved at a higher frequency compared to unlocked movements. Each dot represents the most prevalent conspecific frequency across all movements elicited by a single focal fish. Conspecific frequency was calculated based on a sequence of consecutive movements in the same direction preceding each focal fish movement. The control comparison used a conspecific from a different experiment (Repeated measures ANOVA interaction effect: $p = 0.002$; post-hoc Bonferroni corrected paired t-test in actual data: $p = 9 \times 10^{-5}$, and in control: $p = 0.82$). **F:** Predicted turn angles of focal fish did not differ between locked vs. unlocked (paired t-test, $p = 0.85$) and approach vs. non-approach (paired t-test, $p = 0.99$) movements. **G:** Cumulative distribution of approach probability across all focal fish. 35% of the fish had approach probabilities below chance (gray). **H:** The probability of conspecifics moving rightward relative to the focal fish was higher following a rightward movement compared to a leftward movement, suggesting that conspecifics tend to move in sequences (paired t-test, $p = 1 \times 10^{-24}$). **I:** Probability of focal fish movements occurring at different time points along conspecific sequences (red), indicating a tendency for focal fish to move early in the sequence. Each sequence was divided into four equal bins, representing fractions of its total duration. Chance level probability (black) was calculated by initiating the same number of conspecific sequences at random timings. Shaded areas indicate SEM across all fish. Throughout the figure, $^{***}$ indicates $p < 0.001$, where specified.

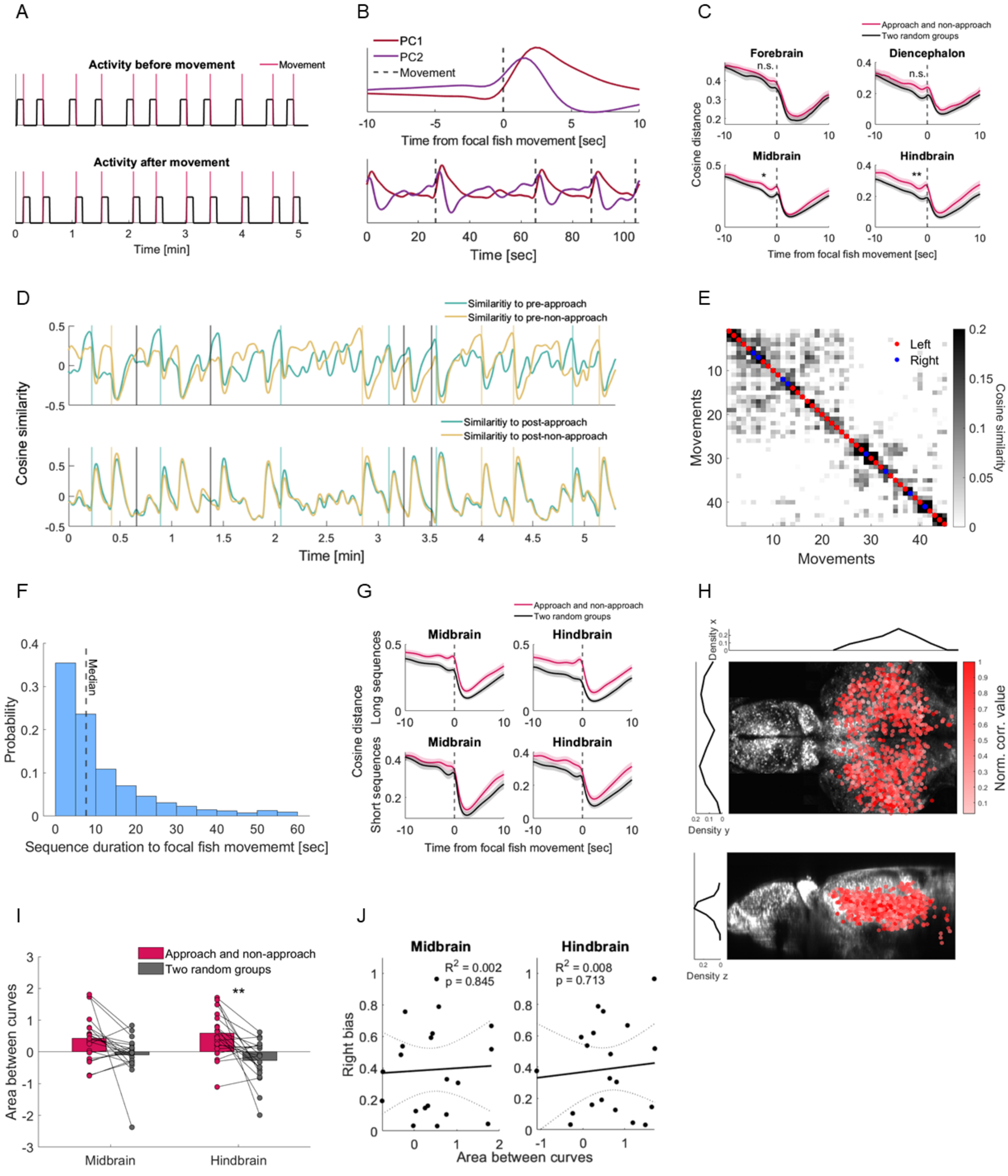

A
Activity before movement
Movement
Activity after movement
Time [min]
B
PC1
PC2
Movement
Time from focal fish movement [sec]
Time [sec]
C
Approach and non-approach
Two random groups
Forebrain
Diencephalon
Midbrain
Hindbrain
n.s.
Cosine distance
Time from focal fish movement [sec]
D
Similaritiy to pre-approach
Similaritiy to pre-non-approach
Similaritiy to post-approach
Similaritiy to post-non-approach
Cosine similarity
Time [min]
E
Left
Right
Movements
Cosine similarity
F
Median
Probability
Sequence duration to focal fish movememt [sec]
G
Approach and non-approach
Two random groups
Midbrain
Hindbrain
Long sequences
Short sequences
Cosine distance
Time from focal fish movement [sec]
H
Density x
Density y
Density z
Norm. corr. value
I
Approach and non-approach
Two random groups
Area between curves
Midbrain
Hindbrain
J
Midbrain
Hindbrain
R2 = 0.002
p = 0.845
R2 = 0.008
p = 0.713
Right bias
Area between curves

**Figure S4: Distinct neural signatures preceding approach movements in the midbrain and hindbrain. A:** Two regressors representing neural activity before (top) and after (bottom) focal fish movements (red lines). **B:** The first two principal components (PCs) of neural activity, averaged across all movements (top) and shown as time series (bottom), revealed two prominent activity profiles: increased neural activity after focal fish movements (first PC) and ramping neural activity before movements (second PC). The dashed black lines indicate movement onsets. **C:** Cosine distance between the mean population neural activity preceding and following approach and non-approach movements compared to the distance between two groups of randomly selected movements. The cosine distance over the five seconds preceding approach and non-approach movements was higher than random, with the extent of this neural distinction varying across brain regions (two-way repeated measures ANOVA, main effect for movement type: $p = 0.003$, interaction effect: $p = 0.048$). This distinction was most pronounced in the midbrain and hindbrain (Post-hoc Bonferroni-corrected paired t-test: forebrain $p = 0.1$, diencephalon $p = 0.17$, midbrain $p = 0.027$, hindbrain: $p = 0.002$). Shaded areas indicate SEM across fish. **D:** Cosine similarity of midbrain population activity to two reference neural patterns preceding (top) and following (bottom) approach and non-approach movements (green and yellow vertical lines, respectively) over five minutes. Top: Population activity before approach movements was more similar to the reference pattern before approach compared to that before non-approach, and vice versa. This difference in similarity disappeared after movement onset. Bottom: Population activity showed the same level of similarity to both references after approach and non-approach movements, indicating that distinct neural patterns were evident only before movement onset. Cosine similarity was computed relative to the mean of a 5-second reference neural pattern, averaged separately for approach and non-approach movements. Black vertical lines indicate movements with small predicted turn angles, which were not classified as approach or non-approach (see Methods). **E:** The similarity matrix shown in Figure 3E, labeled by turning direction of each movement (right: blue, left: red), indicating that the approach and non-approach clusters were independent of turning direction. **F:** Duration of conspecific sequences preceding movements of all focal fish. The median sequence duration is marked by a black dashed line. **G:** Cosine distances of mean population vectors in the midbrain (left) and hindbrain (right) across fish for long (top) and short (bottom) conspecific sequences, showing that approach vs. non-approach distinctions emerged earlier for long sequences. The difference between the two curves (red and black) averaged over 5-10 seconds before movement onset was larger for long compared to short sequences (two-way repeated measures ANOVA, main effect for sequence duration: $p = 0.009$). For each fish, short and long sequences were divided according to the median sequence duration. Shaded areas indicate SEM across fish. **H:** Neurons in the midbrain and hindbrain that most contributed to the cosine distance between approach and non-approach movements (top 100 neurons per region in a single fish, across 6 registered fish) were projected onto a reference brain and were spatially dispersed. Shades of red indicate the strength of correlation. Top and side views are shown at the top and bottom, respectively. **I:** The area between the curves (marked in gray in Figure 3D), computed across fish, was significantly greater than chance in the hindbrain and approached significance in the midbrain (Bonferroni-corrected paired t-test, midbrain: $p = 0.085$, hindbrain: $p = 0.004$). Considerable variability was observed in this area across individuals. **J:** No correlation was found between the area between the curves and movement bias to the right, indicating that the relation between this area and approach probability (Figure 3H) was independent of turning direction. Right bias was calculated as the difference in the fraction of right turns between approach and non-approach movements. Throughout the figure, $^*$ and $^{**}$ indicate $p < 0.05$ and $p < 0.01$, respectively, where specified.

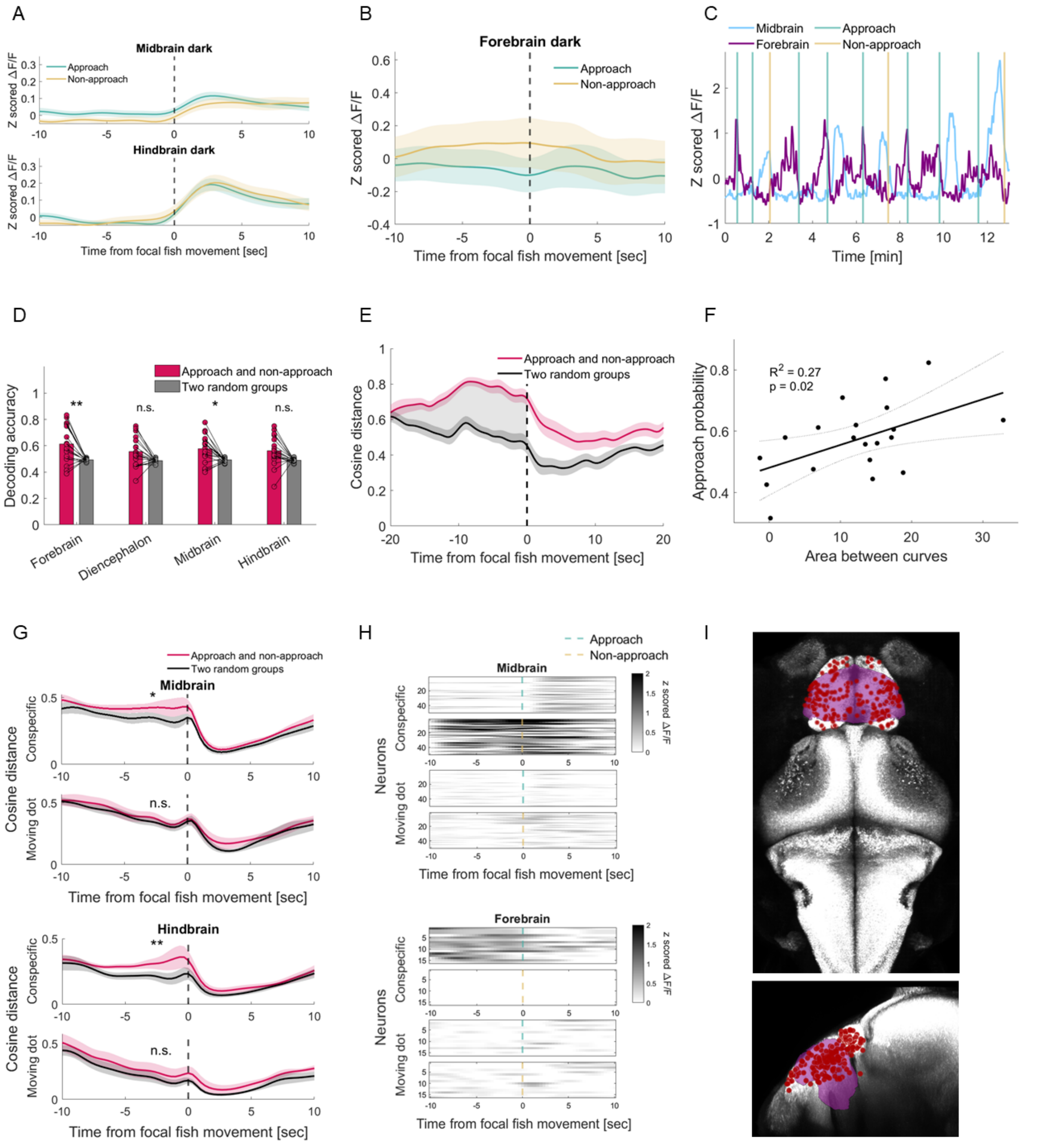
A
Midbrain dark
Approach
Non-approach
Z scored ΔF/F
Hindbrain dark
Time from focal fish movement [sec]
B
Forebrain dark
Approach
Non-approach
Z scored ΔF/F
Time from focal fish movement [sec]
C
Midbrain
Forebrain
Approach
Non-approach
Z scored ΔF/F
Time [min]
D
Approach and non-approach
Two random groups
Decoding accuracy
**
n.s.
*
n.s.
Forebrain
Diencephalon
Midbrain
Hindbrain
E
Approach and non-approach
Two random groups
Cosine distance
Time from focal fish movement [sec]
F
R² = 0.27
p = 0.02
Approach probability
Area between curves
G
Approach and non-approach
Two random groups
Midbrain
Cosine distance
Conspecific
Moving dot
Time from focal fish movement [sec]
Hindbrain
Cosine distance
Conspecific
Moving dot
Time from focal fish movement [sec]
H
Approach
Non-approach
Midbrain
Neurons
Conspecific
Moving dot
z scored ΔF/F
Time from focal fish movement [sec]
Forebrain
Neurons
Conspecific
Moving dot
z scored ΔF/F
Time from focal fish movement [sec]
I

**Figure S5: Forebrain neural dynamics within a distributed network mediate socially specific approach. A:** The distinct neural activity patterns in the midbrain and hindbrain observed before approach and non-approach movements disappeared in the dark, suggesting that these patterns are specific to the observed visual cue. This analysis was based on the same neurons from each focal fish as in Figure 4B. **B:** The distinct forebrain activity patterns observed before approach and non-approach movements were abolished in the dark, indicating that these patterns are specific to the observed visual cue. This analysis was based on the same neurons from each focal fish as in Figure 4E. **C:** Coordinated activity patterns in the midbrain and forebrain reveal distinct and opposing neural dynamics preceding individual approach and non-approach movements. Forebrain activity increased before approach movements, while midbrain activity increased before non-approach movements, with both returning to baseline afterward. Activity patterns were computed by averaging the 14 most distinguishing midbrain neurons (approach vs. non-approach) and the top 14 forebrain neurons most correlated with the approach regressor in an example fish. **D:** Decoding accuracy of the upcoming movement identity based on neurons highly correlated with the approach regressor (Figure 4D, top). Decoding performance was highest in the forebrain compared to other brain regions (Bonferroni-corrected paired t-test, forebrain: $p = 0.007$, Cohen's $d = 1.22$; diencephalon: $p = 0.1$, $d = 0.83$; midbrain: $p = 0.01$, $d = 1.13$; hindbrain: $p = 0.07$, $d = 0.89$). **E:** Cosine distance dynamics between mean population vectors of approach and non-approach movements (red curve), derived from forebrain neurons highly correlated with the approach regressor, compared to a random division of movements (black curve). Red and dark gray shaded areas indicate SEM across fish. The gray shaded area indicates the area between the curves before movement onset. **F:** The neural distinction in forebrain activity before the movement (panel B) accounted for inter-individual variability in approach probability. The solid black line indicates the regression fit, with black dashed lines marking the 95% confidence bounds. **G:** Cosine distance dynamics of mean population vectors in the midbrain (top) and the hindbrain (bottom), using the same neurons from focal fish interacting with a real conspecific or with a synthetic projected dot. The difference between the curves (red and black) averaged over five seconds before movement onset was larger when focal fish interacted with conspecifics (two-way repeated measures ANOVA, main effect for stimulus type: $p = 0.015$). Distinct neural processes for approach and non-approach movements were only present in social interaction (post-hoc Bonfferoni-corrected one-sided paired t-test, midbrain conspecific: $p = 0.039$, hindbrain conspecific: $p = 0.004$, midbrain dot: $p = 0.47$, hindbrain dot: $p = 0.41$). The dashed black lines at time zero represent movement onset. Shaded areas indicate SEM across fish. **H:** Activity profiles of midbrain (top) and forebrain (bottom) neurons that most distinguished approach from non-approach movements in an example fish. When fish observed a real conspecific, decreased midbrain and increased forebrain activity were evident before approach movements, and vice versa before non-approach movements. However, when fish viewed a moving dot, no difference in activity was observed between approach and non-approach movements, suggesting that these neural patterns are specifically related to the social cue. Approach and non-approach movements are marked with dashed green and yellow lines, respectively. **I:** Highly correlated forebrain neurons (216 neurons, 6 fish) were projected onto a 6 dpf reference atlas brain[2], showing a dense concentration in the pallium (purple area). Top and side views are shown at the top and bottom, respectively. Throughout the figure, $^*$ and $^{**}$ indicate $p < 0.05$ and $p < 0.01$, respectively, where specified.
